# A time- and space-resolved nuclear receptor atlas in mouse liver

**DOI:** 10.1101/2023.01.24.525180

**Authors:** Francesco Paolo Zummo, Alexandre Berthier, Céline Gheeraert, Manjula Vinod, Marie Bobowski-Gérard, Olivier Molendi-Coste, Laurent Pineau, Matthieu Jung, Loic Guille, Julie Chevalier-Dubois, David Dombrowicz, Bart Staels, Jérôme Eeckhoute, Philippe Lefebvre

## Abstract

The unique functional versatility of the liver is paramount for organismal homeostasis. Both liver development and adult functions are controlled by tightly regulated transcription factor networks, within which nuclear receptors regulate essential functions of parenchymal and non-parenchymal cells. Acting as transcription factors sensitive to extracellular cues such as steroidal hormones, lipid metabolites, xenobiotics… and modulated by intracellular signaling pathways, nuclear receptors orchestrate many aspects of hepatic physiology. While liver functional zonation and adaptability to fluctuating conditions are known to rely on a sophisticated cellular architecture, a comprehensive knowledge of nuclear receptor functions in the different liver cell types is still lacking. As a first step toward the accurate mapping of nuclear receptor functions in mouse liver, we characterized their levels of expression in whole liver as a function of time and diet, and explored nuclear receptor isoform expression in hepatocytes, cholangiocytes, Kupffer cells, hepatic stellate cells and liver sinusoidal cells. In addition, we leveraged liver single cell RNAseq studies to provide here an up-to-date compendium of nuclear receptor expression in mouse liver in space and time.

## INTRODUCTION

The liver is central to metabolism by coping with qualitatively and quantitatively fluctuating dietary intakes and it stores, packages and reroutes metabolic intermediates to other tissues. The liver also exerts other crucial functions such as detoxification, bile acid synthesis, immune and inflammatory responses and hemostasis. This versatility relies on precisely timed and spatially orchestrated activities of several resident and nonresident cell types which communicate intensively to achieve organ and whole body homeostasis. Within the functional unit of the liver, the hepatic lobule, the above-mentioned biological processes take place into several resident, highly connected cell types [hepatocytes (HC), cholangiocytes (CH), liver sinusoidal endothelial cells (LSEC), stellate cells (HSC) and Kupffer cells (KC)] which are functionally specialized. An additional layer of sophistication is the functional zonation of liver functions, which adapts to the centripetal blood, nutrients and oxygen flow and centrifugal bile circulation (Nagy et al., 2020).

As sensors of the environment through their ability to bind hormones, metabolic intermediates or xenobiotics, nuclear receptors (NRs) are essential relays of metabolic and endocrine signals regulating transcriptional networks in hepatic cells (Soccio, 2020). NR structure allows them to act directly or indirectly as transcriptional regulators (Weikum et al., 2018) and to integrate cues from extracellularly activated signaling pathways (Berrabah et al., 2011). Decade-long research efforts have established that NRs act in a tissue-specific manner through multiple mechanisms ranging from intracellular ligand activation to transcriptional coregulator combinatorial assembly on DNA-bound NR. This notion of a specific activity as a function of the site of expression is likely to be extended to distinct cellular populations within a given tissue, but technical hurdles related to single-cell approaches have to be solved prior to get a full appreciation of NR activity in a specific cell type.

In this respect, the liver is an optimal model to unravel mechanistic aspects of NR actions in cellular subpopulations, as single-cell approaches have paved the way to building a functional atlas of the liver lobule. Based on a thorough knowledge of liver histology, these transcriptomic analysis have partially established the zonation profile of gene expression in mouse and human livers (Payen et al., 2021, Droin et al., 2021, Aizarani et al., 2019, Dobie et al., 2019, Ben-Moshe et al., 2019, Ben-Moshe and Itzkovitz, 2019). However, quantitative and qualitative assessments of NR expression in the liver are scarce (Li et al., 2013, Gonzalez-Sanchez et al., 2017) but needed to fully appreciate their functional diversity and to leverage this knowledge to define innovative therapeutic strategies. Indeed, NR functions have been mostly defined in a hepatocyte background but their expression territory is more diverse. For example, NUR77 encoded by the *Nr4a1* gene is known to modulate hepatic glucose and lipid metabolism (Pols et al., 2008, Pei et al., 2006) and liver regeneration (Hu et al., 2014), but is substantially expressed in CHs, LSECs and KCs from C57Bl/6 mouse liver (Gonzalez-Sanchez et al., 2017, Li et al., 2013) in which its functions are poorly characterized. Similarly, the physiology of the glucocorticoid receptor (GR), has historically been heavily characterized for its role in metabolic regulations (Praestholm et al., 2020), but whose expression in all other liver resident cell types is far from negligible (Gonzalez-Sanchez et al., 2017, Li et al., 2013).

Here, we have leveraged our different transcriptomic studies on bulk and purified liver cells to provide a thorough view of NR isoforms expression in simultaneously isolated parenchymal and non-parenchymal cell populations. Zonation of NR expression was also compiled from published single cell RNAseq studies (Halpern et al., 2018, Bahar Halpern et al., 2017, Su et al., 2021, Dobie et al., 2019) to allow for a refined appreciation of possible physiological NR functions

## MATERIAL AND METHODS

### Animal experimentation

All experiments were approved by the Comité d’Ethique en Expérimentation Animale du Nord-Pas de Calais CEEA75 in compliance with European Union regulations. To eliminate sex as a confounder, only male mice were used throughout this study. C57BL6/J wild-type male mice (12-17 weeks) were purchased from Charles River Laboratories and housed in a temperature-controlled environment (23-25°C) with a 12h/12h light-dark cycle, ZT0 being lights-on. Mice had either free access to water and to a standard chow diet [Safe Diet A04)(“AdLib(itum)” conditions] or access to food was restricted to the active period for 2 weeks prior to euthanasia (12 hours from ZT12 to ZT24)[“T(ime)-R(estricted) F(eeding) condition].

### Multistep isolation of mouse parenchymal and non-parenchymal liver cells

This protocol was optimized to isolate simultaneously hepatocytes (HCs), Kupffer cells (KC), hepatic stellate cells (HSCs), liver sinusoidal cells (LSECs)and cholangiocytes (CHs) from a single liver to obtain sufficient amounts of cells for transcription studies. Livers were obtained at ZT3 from an ad libitum-fed mouse.

#### Cell isolation

Mice (C57Bl6/J male, 12-17-week-old, Charles River) were euthanized by cervical dislocation and liver was perfused through the vena cava. After a first perfusion with Wash buffer [25mM Hepes, pH7.4, 4mM EGTA in 1x Hanks’ Balanced Salt solution (HBSS, Gibco-ThermoFisher #14170)] at 37°C until discoloration of the liver, a second perfusion (≈ 50mL) was performed with Dissociation buffer (25mM Hepes, pH7.4, 1mM CaCl_2_ in 1xHBSS) supplemented with collagenase (type IV, Sigma #C5138)(100U/mL) at 37°C. The liver was then removed and dissociated in a Petri dish. The cell solution was filtered through a 70μm cell filter and centrifuged for 2 min at 50 x G to collect HCs. HC pellets were washed once in 45mL Wash buffer, centrifuged for 2 min at 50 x G and resuspended in FACS buffer [1x phosphate-buffered saline (PBS), pH 7.4, 0.5% bovine serum albumin] supplemented with RNAsin (1:1000, Promega, #N2511) for FACS purification. The HC supernatants from the first centrifugation were collected and spun again for 2 min. at 50 x G to remove remaining HCs from the NPC fraction. NPC cells were then resuspended in Dissociation buffer. Seventy-five % of this preparation (NPC75) was added to the non-digested liver (recovered from the initial 70 μm filtration), centrifuged for 5 min at 580 x G and incubated with collagenase (100 U/mL), and 0.5 mg/mL pronase (Sigma-Aldrich, #10165921001) and 10 μg/mL DNAseI (DNase I grade II, from bovine pancreas, Sigma-Aldrich, # 10104159001) for 20 min at 37°C. The remaining 25% (NPC25) was further digested with collagenase alone for 10 min at 37°C. Cellular preparations were filtered through a 70 μm filter and centrifuged for 5 min at 580 x G in Wash buffer at 37°C. Both NPC fractions are resuspended in 1x FACS buffer supplemented with RNAsin and kept on ice before cell labelling.

#### Cell labelling

All steps from now on were performed at 4°C in light-protected conditions. Cellular fractions were spun down and resuspended in 1 mL red blood cell lysis buffer (155mM NH_4_Cl, 10mM NaHCO_3_, 0.127mM EDTA, pH7.4) for 4 min before adding 1 ml PBS + 0.5% BSA to stop lysis. Two to 3×10^6^ cells were dispatched per tube, centrifuged (5 min, 600 x G, 4°C) and resuspended in 1x PBS, 0.5% Zombie Green (Biolegend, #BLE423112), 1:1000 RNAsin (Promega, #N2511,) for 10 min. After centrifugation, cells were suspended in mouse BD FcBlock (1:200, Blocking anti-CD16/32, Becton-Dickinson #BD 553142), incubated for 15 min on ice and the antibody mix (see below) added for 20 min. Cells were washed twice in 1x FACS buffer and sorted.

#### Cell sorting

Cells were sorted using a BD INFLUX v7 cell sorter (BDBiosciences) driven by the BD FACS Sortware. Compensation particles were from Becton-Dickinson (BD™ CompBeads Compensation Particles Anti-Rat/Hamster Ig, κ Set, # 51-90-9000949). Fluorochrome-coupled antibodies (BioLegend) targeted CD31-BV421 (#BLE102424), CD45-BV510 (#BLE103138), CD326-CF594 (#BLE118236) F4/80-PE-Cy7 (#BLE123114) CD146-APC (#BLE134712) MHCII-AF700-(#BLE107622) CD11b-APC-Cy7 (#BLE101226). Anti-CLEC4F was from R&D (#MAB2784) and coupled to CF568 using the Mix-n-Stain CF568 Antibody Labeling Kit (Biotium, #BTM92235). All antibodies were used at 1:100 dilution. HCs sorting was performed using a 200 μm nozzle and the following settings: pressure 3.7 psi, drop frequency 6.30 kHz, piezo amplitude 4.1, sample fluid pressure was adapted to reach a maximum events rate of 1 000 events/sec. HCs were selected as viable large cells as visualized on FSC/SSC dotplot, and subsequently gated on singlets before sorting (Supplementary Figure 1).

Non parenchymal cells were sorted as follows: the INFLUX cell sorter was equipped with a 86 μm nozzle and tuned at a pressure of 24.7 psi, a drop frequency of 48,25 kHz, a piezo amplitude of 6.7 and sample fluid pressure was adapted to reach a maximum events rate of 10 000 events/sec. NPCs fractions (both NPC75 and NPC25, Supplementary Figure 1B) were gated for viable singlet cells as visualized on FSC/SSC and FSC-W/FSC-A dotplots, respectively and live cells were then selected as “Zombie Green low” events. HSCs were selected as UV^+^ granular cells out of the NPC75 fraction, taking advantage of UV light excitation of retinol and retinoic acid contained in HSCs granules (Mederacke et al., 2015). A “non-small HSCs” gating was applied in order to avoid sorting of degranulated or damaged HSCs. LSECs were selected as UV^-^ CD45^-^ CD146^hi^ events of the NPC25 fraction. CHs were selected as UV^-^ CD45^-^ CD146^low^ CD326^+^ events of the NPC75 fraction. KCs were selected as UV^-^ CD45^+^ F4/80^+^ CLEC4F^+^ events of the NPC75 fraction, and a “non-small KCs” gating was applied in order to avoid sorting of immature or damaged KCs.

Sorted viable HCs were collected in 1 mL RNAlater (ThermoFisher, # 10564445) while sorted viable NPCs were collected in lysis buffer and further processed for RNA extraction. Cytometry data were analyzed using FlowJo v10.5.3 (FlowJo, LLC).

### Immunofluorescence on sorted liver cells

Cell preparations were deposited on a glass slide using a Cytospin 4 (ThermoScientific). Cells were fixed in 4% paraformaldehyde for 10 min and washed twice in 1x Phosphate-Buffered Saline (PBS). After blocking with 10% normal goat serum in 1x Tris-buffered saline (TBS) for 1 hour at room temperature (RT), slides were incubated with the primary antibody for 1 hour at RT or overnight at 4°C. After 3 washes in 1x TBS, secondary antibodies were added at the indicated dilution in 1x TBS for 1 hour at RT. Slides were washed as above and prepared for microscopy.

Primary antibodies used were: Anti-KRT18 (C-04, Abcam, #ab668)(dilution 1:100), anti-DESMIN (Y66, Abcam, #ab32362)(dilution 1:50), anti-CLEC4F (ThermoFisher, # MA5-24113)(dilution 1:100), anti-VECAD (ThermoFisher, # 36-1900)(dilution 1:50), anti-KRT19 (EP1580Y, Abcam, # ab52625)(dilution 1:100). Secondary antibodies (ThermoFisher Scientific) were used at 1:100 dilution and were goat anti-mouse AF568 (A-11004), donkey anti-rabbit AF488 (A-21206), donkey anti-mouse AF555 (A-31570) and goat anti-rat AF488 (A-11006).

### RNA extraction and RT-qPCR

RNA was extracted using the Macherey-Nagel™ Mini kit Nucleospin™ (Macherey-Nagel, # 872061) or Qiagen RNeasy micro kit (Qiagen, # 74004), depending on abundance of cell preparations, following the manufacturer’s instructions. RNA concentration and purity were assessed using a Nanodrop One device (ThermoFisher Scientific) or a Qubit fluorometer (ThermoFisher Scientific) and a Qubit RNA HS Assay kit (ThermoFisher Scientific, # Q32852), while RNA integrity was analyzed on a Bionanalyzer 2100 (Agilent). RNA preparations with RIN<6.0 were discarded. RNAs were reverse-transcribed using random primers and the High Capacity cDNA Reverse Transcription Kit (ThermoFisher/Applied Biosystems, # 4368814). Quantitative PCR was performed in technical triplicates from at least 3 independent biological samples using the SYBR green Brilliant II fast kit (Agilent Technologies) on an Mx3005p apparatus (Agilent Technologies) or a QuantStudio 3 (Applied Biosystems). Expression values obtained from mRNA levels normalized to *Rps28* (ribosomal protein S28) and *Rplp0* (acidic ribosomal phosphoprotein P0) mRNA levels and were used to calculate fold changes using the cycle threshold (2^−Δ Δ Ct^) method (Schmittgen and Livak, 2008). Primer sequences are listed in Supplementary Table 1.

### Affymetrix array analysis

#### RNA processing and array hybridization

Gene expression from whole mouse liver (n=3) was analyzed with Affymetrix GeneChip MoGene 2.0 ST arrays after RNA amplification, sscDNA labeling and purification. Briefly, RNA was amplified using the GeneChip™ WT PLUS Reagent Kit (Thermo Fisher Scientific, # 902280), retrotranscribed to sscDNA and labeled using GeneChip™ WT Terminal Labeling Kit (Thermo Fisher Scientific, # 900670), followed by hybridization on the GeneChip Mouse Gene 2.0 ST Array (Affymetrix, # 902118) according to the manufacturer’s instructions.

#### Data processing and analysis

Raw data were processed further using GIANT, a user-friendly tool suite developed in-house for microarray and RNA-seq differential data analysis (Vandel et al., 2020). It consists of modules allowing to perform quality control (QC), Robust Multi-Average method normalization, LIMMA differential analysis, volcano plot and heatmaps. Differential gene expression was analyzed after normalization of signals to the median of all samples, log_2_ transformation and exclusion of the 10^th^ lowest percentile considered as technically unreliable.

### RNA sequencing

#### Library preparation and sequencing

RNA samples were sent to the GenomEast platform for library preparation and sequencing. Briefly, RNA preparations were first depleted from unwanted, abundant transcripts using Ribo-Zero Plus rRNA depletion kit (Illumina, # 20040526). cDNA synthesis, 3’end adenylation, adapter ligation and PCR amplification were performed using the TruSeq Stranded total RNA sample preparation kit (Illumina). DNAs were sequenced using an Illumina HiSeq 4000 sequencer in 50 bp Single-Read following Illumina’s instructions. Image analysis and base calling were performed using RTA 2.7.7 and bcl2fastq 2.17.1.14. Sequencing depth was 75 million reads on average.

#### Data processing and analysis

Reads were preprocessed using cutadapt version 1.10 (Kechin et al., 2017) in order to remove adapter, polyA and low-quality sequences (Phred quality score below 20). Reads shorter than 40 bases were discarded for further analysis. Remaining reads were mapped onto the mm10 assembly of the Mus *musculus* genome using STAR version 2.5.3a (Dobin et al., 2013).

Gene expression quantification was performed from uniquely aligned reads using htseq-count version 0.6.1p1 (Anders et al., 2015), with annotations from Ensembl version 94 and “union” mode. Read counts were normalized across samples with the median-of-ratios method (Anders and Huber, 2010), suitable for inter-sample comparison. Gene expression profiles were compared using the Bioconductor package DESeq2 version 1.16.1 (Love et al., 2014). P-values were adjusted for multiple testing using the Benjamini and Hochberg method (Benjamini and Hochberg, 1995).

#### Splice junctions and isoform detection

Splice junctions were visualized in Integrative Genomics Viewer (IGV, Broad Institute)(Robinson et al., 2011) using bam files of aligned reads and the mm10 gene annotation track. Alignment data were visualized using the Sashimi plot function of IGV.

#### Single cell data extraction

Analysis were carried out using extracted data from available datasets (GSE84490 for HC zonation and GSE108561 for LSEC zonation)(Halpern et al., 2018, Bahar Halpern et al., 2017) or using the ISCEBERG browser which allows analysis and interrogation of single cell RNAseq data (Guille et al., 2022) for HSC zonation [GSE137720 (Dobie et al., 2019)] and LSEC zonation [GSE147581, (Su et al., 2021)].

#### HSCs, GSE137720

Seurat (v 4.0.1) was used to analyze this dataset. According to (Dobie et al., 2019), we filtered out cells expressing < 300 genes and cells expressing > 30 % of mitochondrial genes. Then data were normalized and scaled using NormalizeData and ScaleData functions from the Seurat package. We applied a batch correction using Harmony (v 0.1.0). Then dimensionality reduction was achieved using Uniform Manifold Approximation and Projection (UMAP) to calculate 2D coordinates. SCINA (v 1.2.0) was used to characterize cell populations (fibroblasts, vascular smooth muscle cells, and hepatic stellate cells) according to cell markers defined in (Dobie et al., 2019). Cells distinct from HSCs were filtered out and HSCs located in the periportal or pericentral areas were identified using SCINA (version 1.2.0) based on markers used in the publication.

#### LSECs, GSE147581

Seurat (v 4.0.1) was used to analyze this dataset. According to (Su et al., 2021), cells expressing less than 200 transcripts and more than 20 % of mitochondrial genes were filtered out. Data were normalized and scaled, and Harmony (version 0.1.0) was used to apply a batch correction. UMAP coordinates were calculated and clusterized with findClusters using a resolution of 0.5 to match with the published analysis. Annotation with identified zonation markers was carried out using SCINA (version 1.2.0).

### Statistical analysis

Statistical analysis was performed using GraphPad Prism (v. 9). Data are plotted as the mean ± SEM. At least 3 independent experimental replicates were obtained. Data were determined to have equal variances using the F test. For 2-group comparisons, an unpaired 2-tailed *t*-test with Welch correction was used. For multiple comparisons with one variable, a 1-way ANOVA followed by the Tukey multiple comparison test (each group compared to every other group) was used. Multiple comparisons with more than one variable were carried out using a 2-way ANOVA followed by a Tukey’s multiple comparison test. Cyclical patterns of gene expression were determined using JTK_Cycle (Hughes et al., 2010). Individual expression/level data were plotted in Prism as a time series and non-linearly fitted using the “sine wave with nonzero baseline” function, using a wavelength constraint previously determined by JTK_cycle (in the 20-24 hours range).

### Data visualization

#### Bubbleplots

Bubbleplots were generated in R studio using the ggplot2, plotly, reshape2, rcpp and tidyverse packages. SVG files were modified with Inkscape v1.0 and assembled as figures using CorelDraw2020. The liver trabeculae structure was adapted from a file published in a Public Library of Science journal (Frevert et al., 2005) under the Creative Commons Attribution 4.0 license.

### Data availability

Affymetrix data files (“AdLib” and “TRF” data) are available under the GEO dataset number GSE223360. RNA sequencing data are available under the GEO dataset number GSE222597.

## RESULTS

### Development of a liver cell type multi-step isolation protocol

In order to minimize both technical and biological biases in isolating liver cell populations, we set up a protocol allowing the purification of 5 resident cell populations, i.e. HCs, LSECs, HSCs, KCs and CHs from a single liver. This protocol also allowed the purification of dendritic cells and of neutrophils, which were not considered further in this study. After sacrifice by cervical dislocation to avoid any side effects of anesthetics, the liver was perfused with modified HBSS and dissociated with collagenase IV. To enrich for specific cell populations, aliquots of the digested liver were then processed separately (Figure 1). Dissociated cells were sorted based on size to yield purified HCs, whose amounts routinely exceeded 20 ×10^6^ cells per liver. Further digestion by collagenase and pronase yielded the total non-parenchymal cell (NPC) fraction which was sorted using the indicated combination of antibodies (Figure 1 and Supplemental Figure 1). This yielded per liver variable amounts of HSCs, KCs, LSECs and CHs with numbers routinely exceeding 10^5^ cells per preparation and cell type (Supplemental Figure 2). Cellular homogeneity and purity were assessed by RT-qPCR and immunofluorescence using established cell type-specific cellular markers (Figures 2 and Supplemental Figure 3).

**Figure 1.**
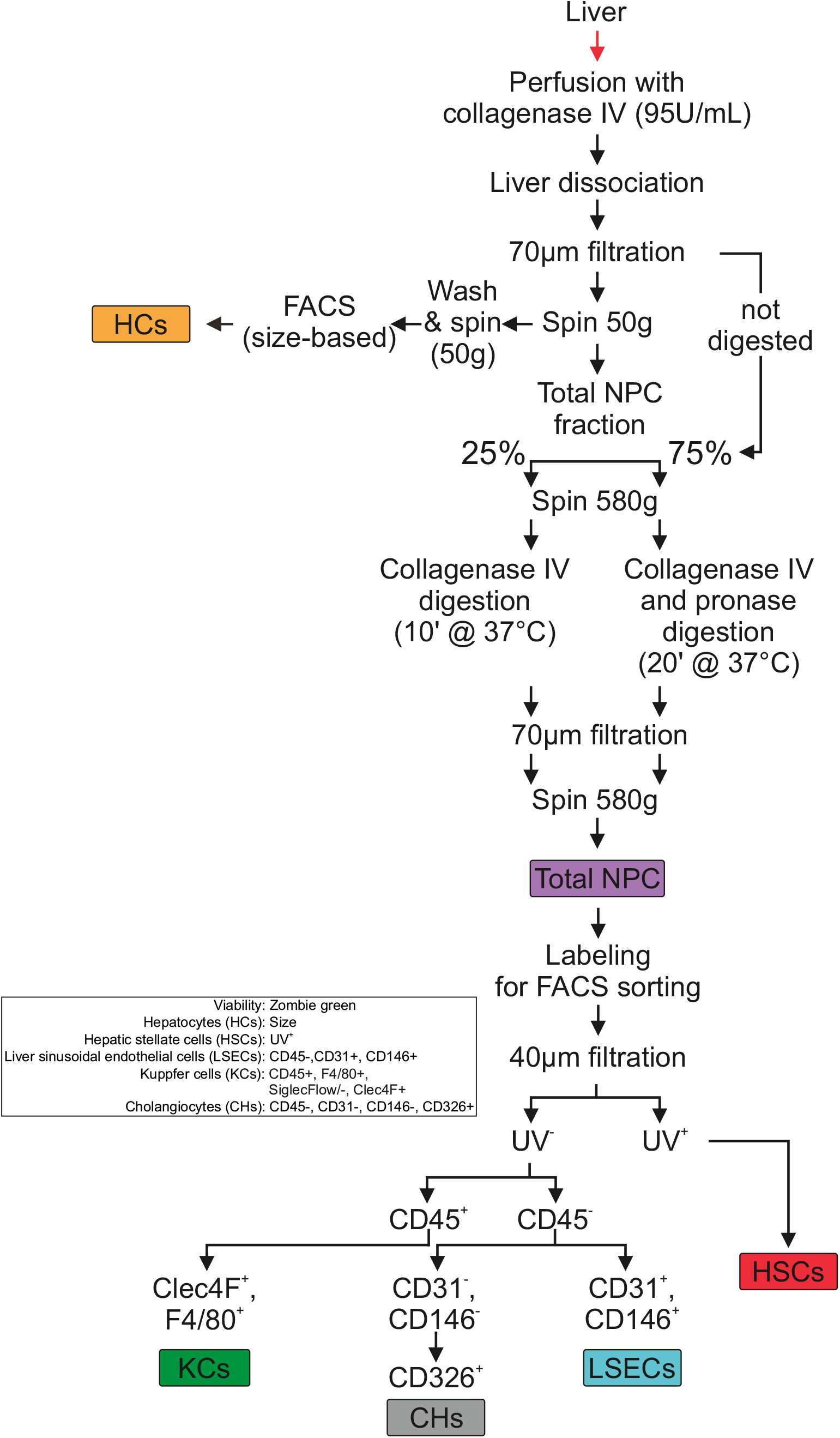
Liver cell type isolation protocol. The sequential steps of liver cell type purification by fluorescence-activated cell sorting (FACS) are shown here. FACS parameters are indicated (box) and details of the complete procedure can be found in the Material & Methods section. HCs: hepatocytes; NPC: nonparenchymal cells; KCs Kupffer cells; CHs: cholangiocytes; LSECs: liver sinusoidal endothelial cells; HSCs: hepatic stellate cells. An example of the FACS output is shown in Supplemental Figure 1.

**Figure 2.**
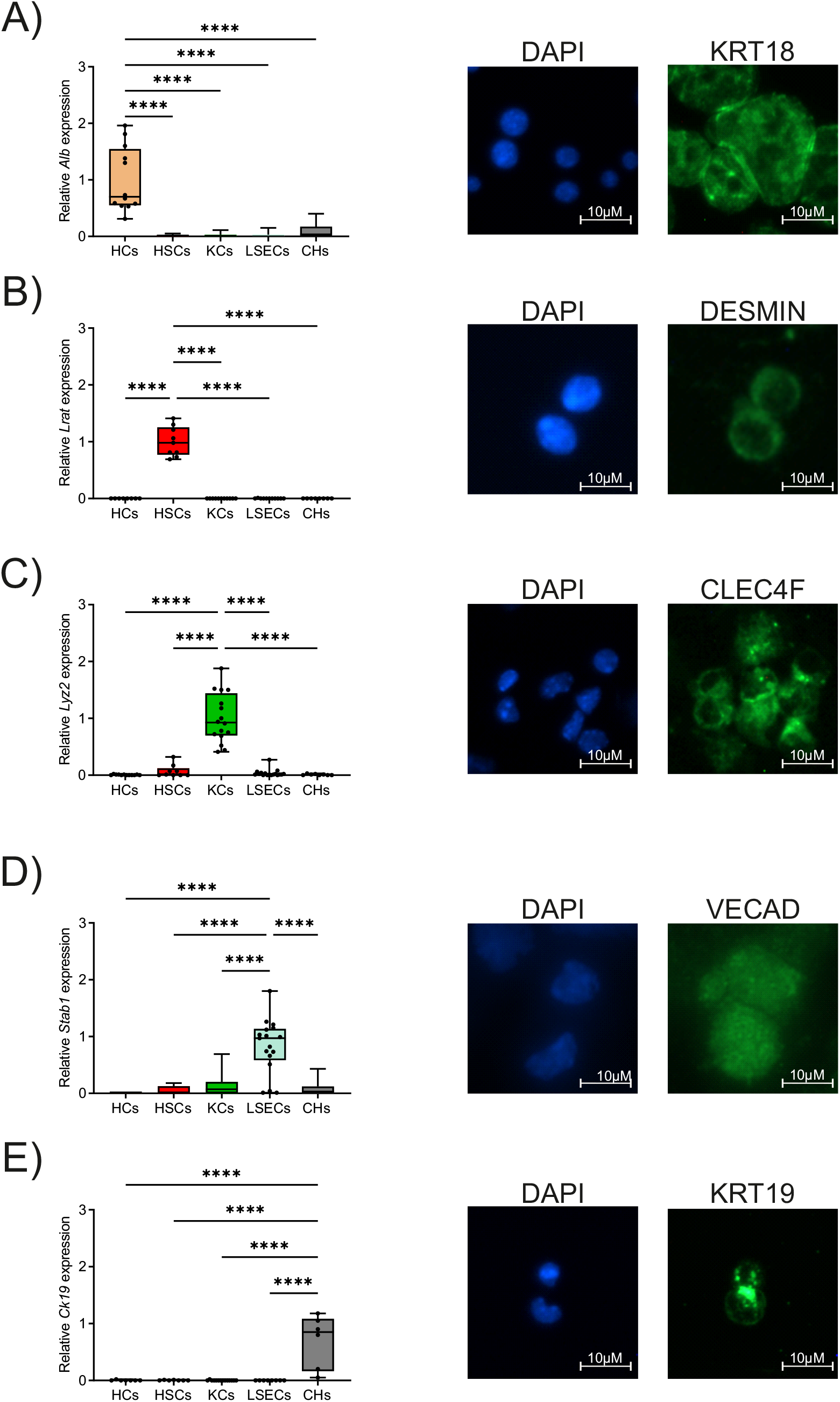
Characterization of purified liver cell types. After FACS-based purification, RNA was extracted and used for RT-qPCR assays (left panels) or cells were deposited on a glass slide using Cytospin centrifugation to be further labelled with the indicated antibodies. A) HCs characterization; B) HSCs characterization; C) KCs characterization; D) LSECs characterization; E) CHs characterization.

**Figure 3.**
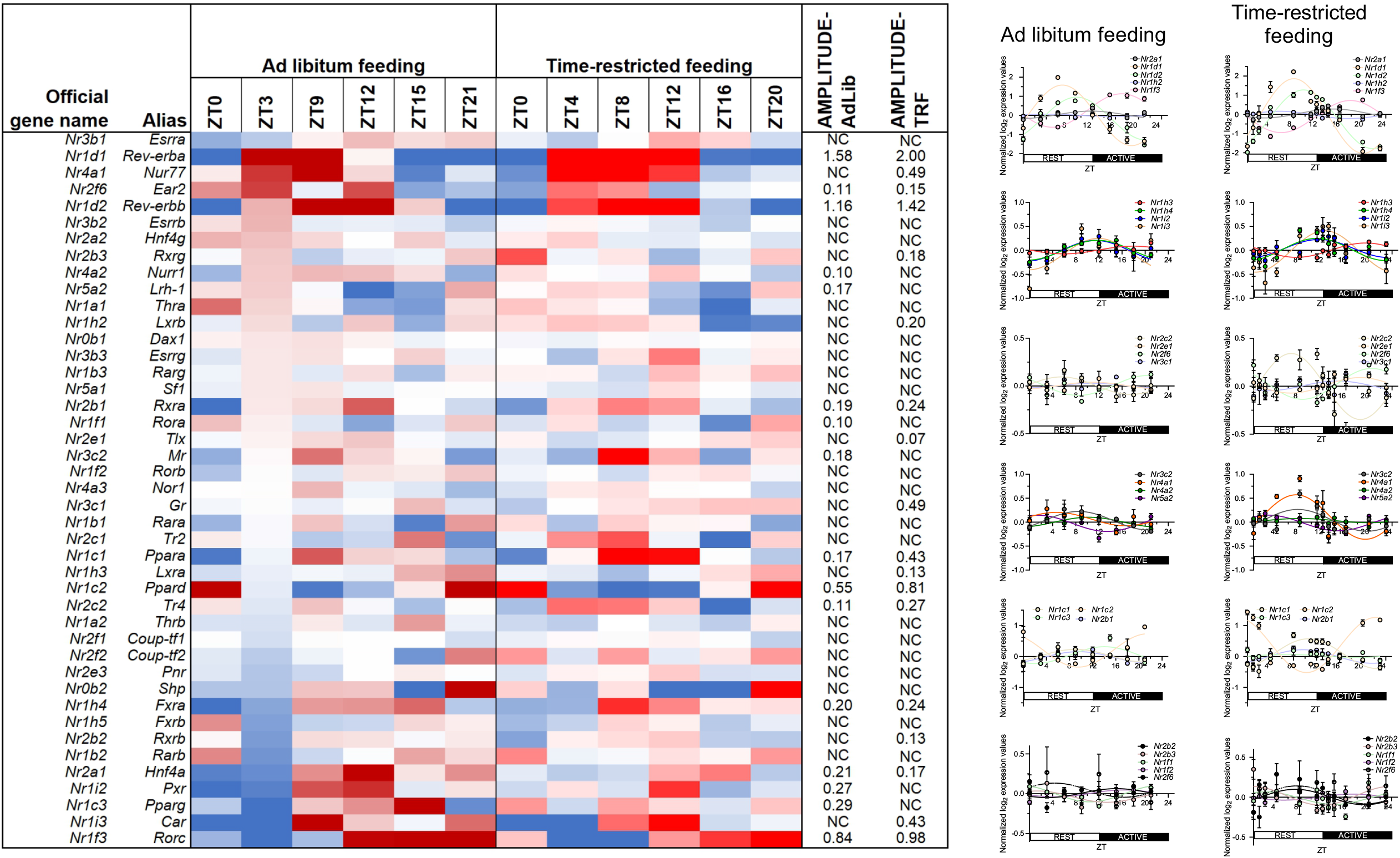
Circadian expression of nuclear receptors. Mice were fed a chow diet either ad libitum or under a time-restricted regimen for 2 weeks. Liver were collected at indicated times (ZT0 being “lights-on) and extracted RNAs were analyzed on Affymetrix arrays (n=3). Data were processed and gene expression values for NRs were extracted and used to generate a heatmap (left panel). Transcriptomic data were also analyzed using the JTK R script to identify genes displaying a cyclic expression. Calculated amplitudes for NR-encoding genes with a 24h cyclicity are indicated (left panel, NC: non-cyclic). Differences in phases and amplitudes were explored by fitting expression data by cosine curves and each point represents the mean ± S.E.M. (n=8). ZT: zeitgeber time.

### Circadian rhythmicity of hepatic nuclear receptors expression

In homeostatic conditions, NR expression may vary not only as a function of nutritional and hormonal cues, but also according to the day-night cycle. To assess whether time-of-the-day is a critical parameter in dictating NR-encoding transcript abundance, we compared gene expression patterns in C57Bl6 male mice liver fed a chow diet either *ad libitum* or under time-restricted feeding. In the latter case, food was available only during the active period (dark period for 12 hours). Transcriptomic data were obtained and analyzed using the JTK package to determine gene expression periodicity and amplitude (Hughes et al., 2010)(Table 2). Several NRs displayed a robust 23 to 24-hours cycle (*Nr1d1*/*Rev-erba, Nr1d2*/*Rev-erbb, Nr1c2*/*Pparb/d* and *Nr1f3*/*Rorg*) while *Nr2f6*/*Ear2, Nr2b1*/*Rxra, Nr1c1*/*Ppara, Nr2c2*/*Tr4, Nr1h4*/*Fxra* and *Nr2a1*/*Hnf4a* cycled similarly albeit with a lesser amplitude (Figure 3). In most cases, a time-restricted feeding regimen increased the amplitude of oscillations, without altering the phase (right panels), in line with a recent report (Deota et al., 2023). More surprisingly, several NRs exhibited condition-specific cycling (ad libitum-fed: *Nr4a2*/*Nurr1, Nr5a2*/*Lrh1, Nr1f1*/*Rora, Nr3c2*/*Mr, Nr1i2*/*Pxr, Nr1c3*/*Pparg*; time-restricted feeding: *Nr2b3*/*Rxrg, Nr1h2*/*Lxrb, Nr3c1*/*Gr, Nr1h3*/*Lxra, Nr2b2*/*Rxrb, Nr1i3*/*Car*). Irrespective of the functional consequences of such oscillations, these differential expression levels should be considered when comparing expression levels of NRs in different conditions. We thus selected ZT3 (3 hours after light-on) as a convenient reference time point to initiate liver cell type isolation from ad-libitum fed mice (Figure 1). In these conditions, *Nr1d1*/*Rev-erba* and *Nr1d2*/*Rev-erbb* reached their zenith, while *Nr1f3*/*Rorg, Nr1c3*/*Pparg* and *Nr1i2*/*Pxr* were at their nadir.

### Nuclear receptor expression in liver cell types

NR-encoding transcripts were quantified in each cell type by RNAseq (Figure 4 and Table 3). Forty-two NRs reached a detectable level of expression (RPKM>10), and each cell type was characterized by a specific NR pattern of expression, with a high level in *Hnf4a, Ear2, Ppara* and *Car* mRNAs being characteristic of hepatocytes. Cholangiocytes exhibited a highly restricted panel of highly expressed NRs, including only *Rora* and *Hnf4g*. LSECs showed high levels in *Rarb* and *g, Coup-tf1&2*, while HSCs were characterized by a high level in *Fxra* and *b, Thra* and *Rara*. Finally, *Rxrb, Nur77, Pparg* and *Lxra* elevated levels were a feature of KCs. On the opposite, hepatocytes were characterized by undetectable levels in *Nor1* and LSECs by the total absence of *Hnf4g*. NRs deemed to be undetectable in any cell type (<10 RPKM) were *Erb*/*Nr3a2, Dax1*/*Nr0b1, Tlx*/*Nr2e1, Pnr*/*Nr2e3, Sf1*/*Nr5a1, Rorb*/*Nr1f2* and *Pr*/*Nr3c3*.

**Figure 4.**
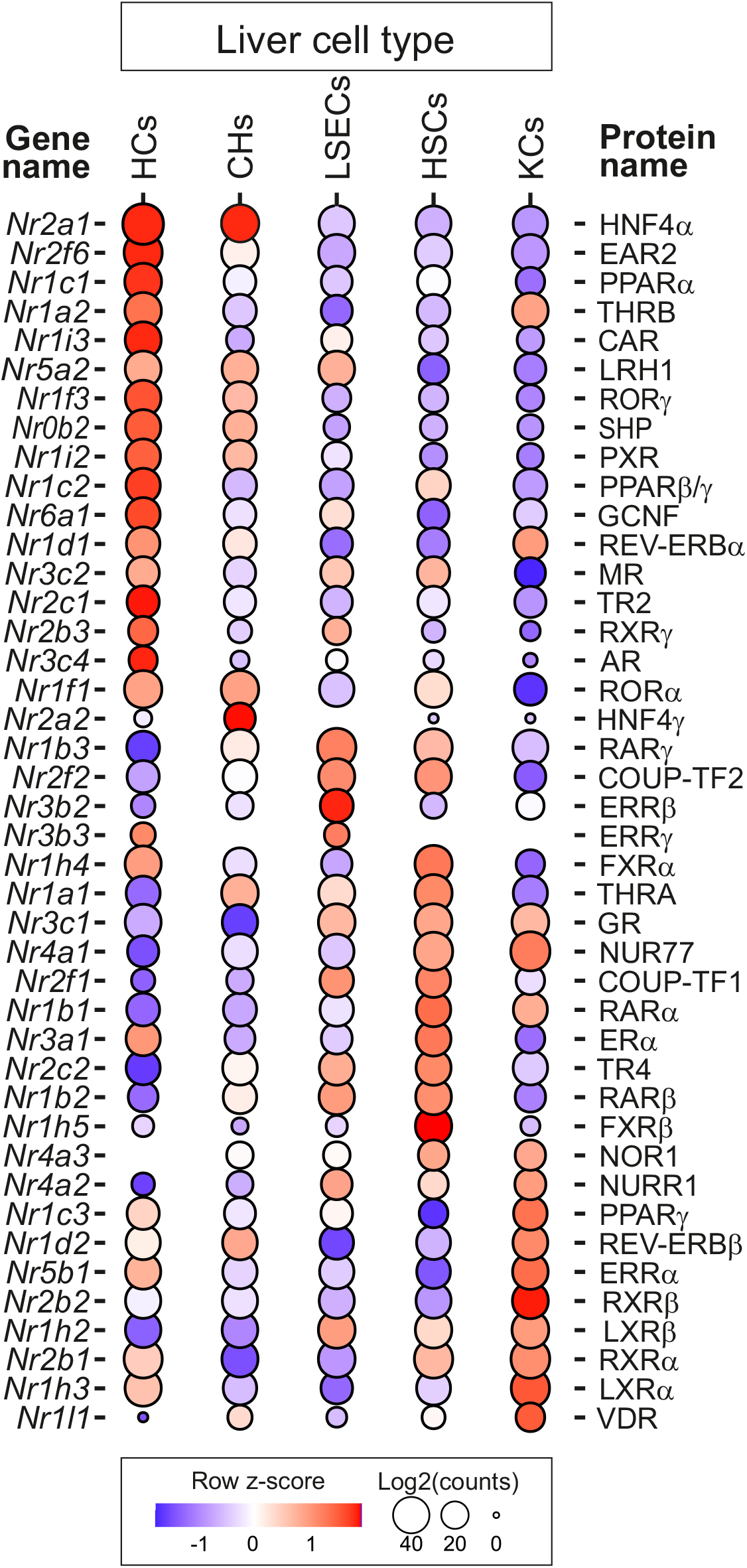
NR expression in purified liver cell types. RNA extracted from each cell type preparation (n=3) was analyzed by single-end 50b RNAseq. After (pre)processing, mapping and normalization by the median-of-ratios method to make counts comparable between samples, log_2_ expression values were used to generate a bubble plot in which row Z-score of TPM (p adj < 0.05) is indicated (genes in red are up-regulated, genes in blue are down-regulated) on a row-by-row basis. The size of the bubble is proportional to the expression level (empty spaces indicate no significant expression).

### Nuclear receptor isoforms expression in liver cell types

Nuclear receptor isoforms play substantially distinct physiological roles, hence determining their expression level is of importance to decipher NR cell-specific functions. NR protein isotypes and their corresponding isoforms were compiled from Uniprot and Protein Ontology databases (UniProt, 2021, Natale et al., 2017), and associated transcripts were searched in RNAseq data using the Sashimi plot function from the Integrative Genome Viewer [IGV, (Katz et al., 2015, Robinson et al., 2011)]. Twenty out of the 42 detected NRs-encoding transcripts can be potentially expressed as distinct isoforms (Figure 5 and Table 3) out of which 11 actually displayed differential expression in the 5 isolated liver cell types. These included transcripts coding for FXRα, PXR, CAR, GCNF, PPARγ, RARα, β and γ, RORα, RXRβ, T3Rα and β, whose expression levels were qualitatively assessed as described and were reported in Figure 5.

**Figure 5.**
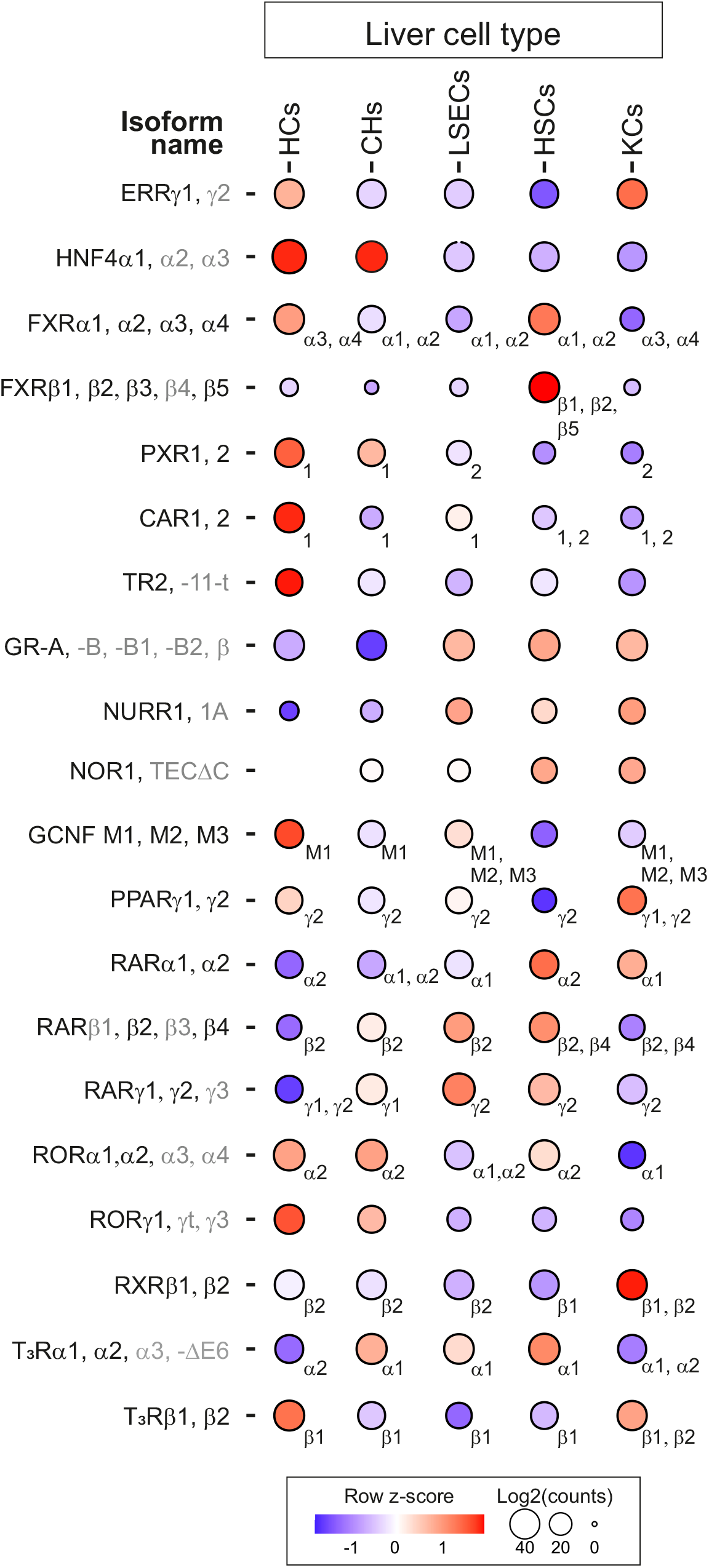
NR isoforms expression in purified liver cell types. Gene expression data were reported form Figure 4 and Uniprot-identified isoforms were indicated. Non-detected isoforms are indicated in gray. (Co)Identified corresponding transcripts are indicated for each cell type and NR.

### Zonation of nuclear receptor expression in hepatocytes, sinusoidal endothelial cells and stellate cells

Functional zonation of the liver is observed along a periportal-pericentral axis and is conditioned by multiple factors such as oxygen, nutrient and morphogen gradients (Panday et al., 2022, Kietzmann, 2017). Specialized functions of liver cell types as a function of their spatial distribution can be inferred from single-cell studies and have been molecularly detailed in recent years. Expression patterns of NR-encoding genes were extracted from published datasets for mouse HCs, LSECs and HSCs (Halpern et al., 2018, Bahar Halpern et al., 2017, Dobie et al., 2019, Su et al., 2021). In HCs, NRs displayed distinct spatial expression patterns)(Figure 6), with PPARα being equally expressed along the pericentral to periportal axis, in agreement with its ability to regulate fatty acid oxidation (predominantly periportal) and ketogenesis (predominantly pericentral). Other NRs also displayed an even gene expression pattern along this axis (*Mr, Rorα* and *γ, Gcnf, Thrβ, Coup-tf2, Shp*), while some had a dominant pericentral localization (*Pparγ, Errβ, Lrh1, Gr, Rxrβ*). Only *Rev-erbβ* and *Erα* were preferentially expressed in the portal area. Mining the transcriptome of LSECs obtained by paired-cell RNAseq (Halpern et al., 2018) defined NR expression in this cell population (Supplemental Figure 4). Thirty-seven NRs were found to have a spatially differential expression, with *Tr2, Lrh1* and *Pparg* being almost exclusively expressed in the pericentral area. Mirroring this pattern, *Pxr, Era* and *Errg* were exclusively detected in the periportal area, while *Coup-tf2, Lxrb, Ppara, Gr, Car, Rxra* and *Errb* were significantly expressed, albeit with variation, in all 4 layers. A comparison with CDH5 (VE-cadherin)-expressing LSEC single-cell transcriptome data brought further elements of comparison, while providing novel information about arterial (portal) and venous (central) LSECs (Supplemental Figure 5). Zonation patterns matched for 50% (18 out 37) NRs, showed minimal discrepancies for 15 and were strikingly different for *Ppara, Erra, Errg* and *Era. Ppara* expression levels, which are in LSECs 4% of that found in HCs, was restricted to arterial-like ECs (Supplemental Figure 5) or present all along the pericentral-periportal axis (Supplemental Figure 4). Along this axis, *Era, Erra* and *Errg* displayed an opposite gradient of expression in these 2 datasets. Cell isolation and identification methods as well as transcript mapping procedures were different in those 2 studies, calling for additional strictly comparative studies to reach a consensual cartography of NRs in LSECs. Of note, our LSEC purification procedure relies on a CD31/CD146 double-positive labeling to obtain highly pure cell preparations, which may have nevertheless selected a particular subpopulation.

**Figure 6.**
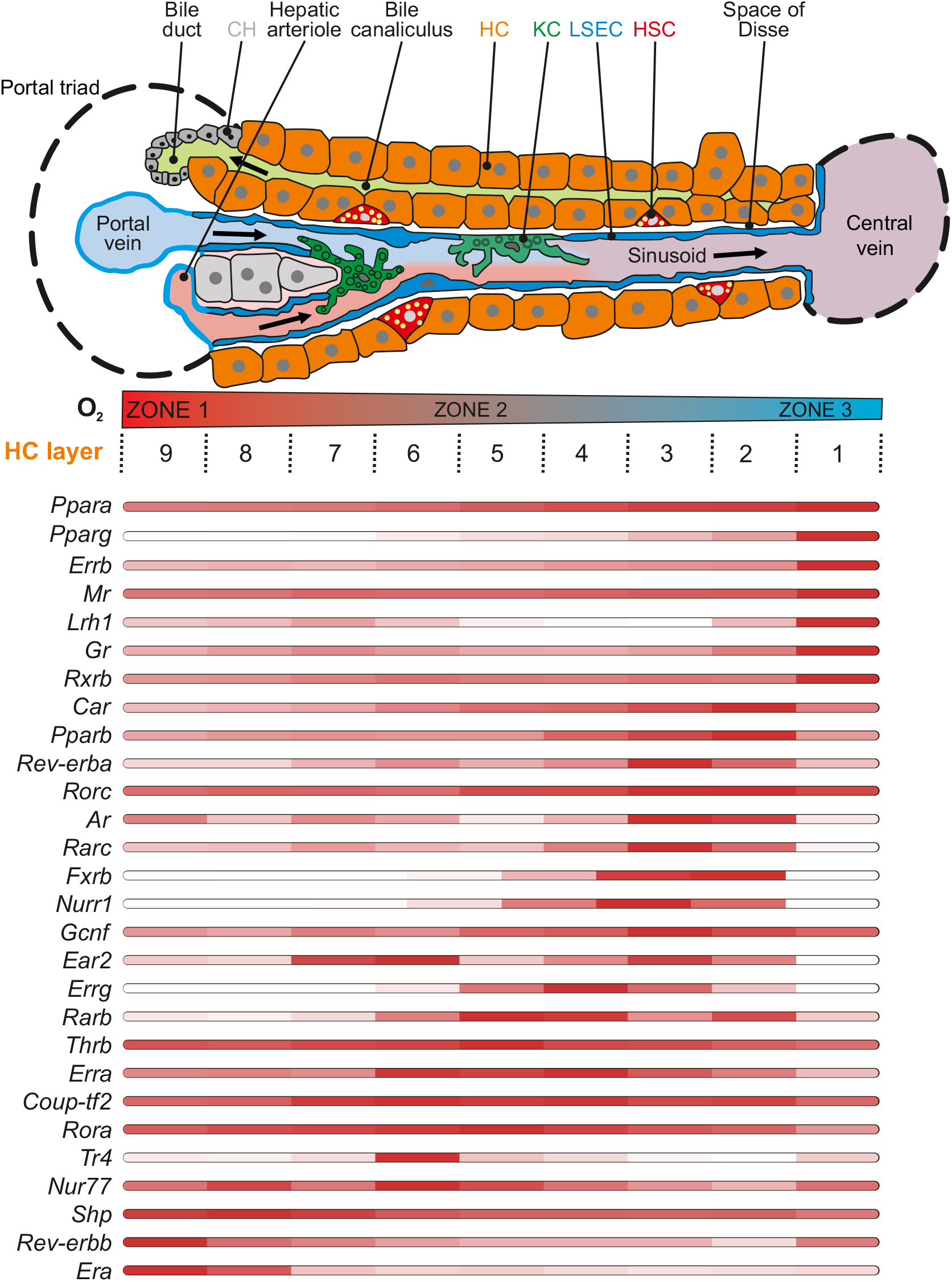
NR zonation in hepatocytes. Upper panel: Schematic organization of a liver trabeculae [(adapted from Wikimedia Commons and initially published in (Frevert et al., 2005)]. Lower panel: expression values for each NR were extracted from (Bahar Halpern et al., 2017) and used to compute an heatmap. Red: High expression, white: low expression. Arrow (right to left) indicates the bile flow, arrows (left to right) indicate the blood flow.

Finally, NR expression was mapped in the 2 identified HSC populations which locate in close vicinity to the periportal (PaHSCs) or of the pericentral (CaHSC) areas (Supplemental Figure 6). With the exception of *Fxra* whose expression was detected in both HSC subpopulations and slightly higher in PaHSCs, the 30 quantified NR-encoding transcripts showed a markedly unbalanced expression between the 2 subpopulations. *Coup-tf2, Gr, Rxra, Rora, Lxrb* and *Ear2* were more preferentially expressed on PaHSCs, whereas *Thra, Nur77, Fxrb*, and *Rara* transcripts were prominently localized in CaHSCs.

## DISCUSSION

Nuclear receptors play critical roles in liver physiology, and establishing a precise spatiotemporal atlas of their expression is mandatory to define their functions. Recent progress in single-cell technologies has shed some light on processes driving liver functional zonation and allowed to map cell-specific expression patterns of NRs, but they still lack sensitivity to identify transcript isoforms in isolated cells. Here we provide a compendium of hepatic NR expression taking into account and minimizing whenever possible technical variability, diet and time-of-the-day influences. By performing a bulk RNAseq analysis of isolated cell types, NR transcript isoforms were also easily identified, as NR protein isoforms are known, at least for a few cases, to bear distinct functional properties. Finally, spatial expression of NRs has been explored by mining single cell transcriptomic datasets, which may provide a mean to ascribe, or rule out, novel functions to NRs.

Our data were compared with others, which were obtained from *ad libitum*-fed C57Bl/6J male (Gonzalez-Sanchez et al., 2017) or female mouse livers collected at non indicated times and using distinct cell isolation and RNA quantification methods. While methodologies were different, these 3 sets of data identified unambiguously NRs which are never detected in any liver cell types (*Dax1, Rorβ, Tlx, Pr* and *Sf1*). Identifying an NR-based cell type signature requires to exclude NRs with oscillating levels of transcripts and strongly sensitive to feeding conditions. Since ad libitum feeding is most commonly used in animal facilities, we used this experimental condition as a reference and could identify *Rev-erba, Rev-erbb, Rorg, Pxr* and *Car, Fxra* and *Pparb/d* as genes with markedly oscillating transcripts along the day-night cycle. Of note, *Nur77*- and *Tr4*-encoded transcripts gained strong cyclicity in time-restricted fed mice. An NR consensus signature characteristic of HC could be defined which identified, when integrating isoform expression patterns, *Hnf4a1, Ppara* and *Thrb1* as being overwhelmingly expressed in this cell type, with 10x expression ratios when compared to CHs, HSCs, LSECs or KCs. In CHs, *Rora2* and *Gcnf M2* isoforms expression were markedly higher than in other cell types (x5 to x10). In LSECs, both *Coup-tf2* and *Rarg2* displayed highest levels of expression, while *Era* and *Fxra1* and *a2* expression were hallmarks of HSCs. Finally, KCs displayed highest levels in *Erra, Lxra* and *Nur77*. Our data thus bring additional information about NR isoform expression, which are in most cases in good agreement with previous reports for NRs displaying high to moderate expression levels. Some minor discrepancies were observed for NRs displaying low expression levels, which are reported as not expressed in PCR-based investigations, but nevertheless detected in the more sensitive RNAseq assay.

Nineteen NRs detected in our study have referenced protein isoforms, a number likely to be vastly underestimated (Annalora et al., 2020). Therefore splicing events could dramatically extend the functional repertoire of NRs, as described for “metabolic NRs” (Mukha et al., 2021). While our study was not designed to formally quantify all alternative transcripts for each NRs expressed in each liver cell type, 13 NR-encoding genes were actually expressed as different isoforms. They included *Fxra, Fxrb, Pxr, Car, Gcnf, Pparg, Rara, b* and *g, Rora, Rxrb, T3Ra* and *b*. Various *scenarii* were observed with respect to isoform expression profiles. A single isoform could be detected per cell type (*Pxr*) or a single or a mix of isoforms were identified (*Fxra, Fxrb, Car, Gcnf, Pparg, Rara, Rarb, Rarg, Rora, Rxrb*). In most cases, isoform expression is known to result from alternative promoter usage (*Fxra1, a2* vs *Fxra3, a4*; *Pparg1* vs *Pparg2*; *Rara1* vs *Rara2*; *Rarg1* vs *RARg2*; *Rxrb1* vs *Rxrb2*) and in the remaining cases (*Rarb2* vs *Rarb4* and *Rora1* vs *Rora2*) from alternative splicing. While the specific functions of NR isoforms has not been studied in great details with a few exceptions (FXR, PPARγ), reports generally point at distinct transcriptional activities and tissue-specific expression of these variants (Mukha et al., 2021). This knowledge has to be refined by investigating the role of NR isoforms in liver cell subpopulations, which exert distinct roles that still remain to be explored. In this respect, the subtissular repartition of FXR, which in contrast to 2 reports (Verbeke et al., 2014, Fickert et al., 2009), we and others (Gonzalez-Sanchez et al., 2017, Garrido et al., 2021) found to be highest expressed in HSCs and less abundantly in HCs, should be refined in light of isoform expression territories. FXR isoform functions have indeed been studied by elegant approaches solely in a hepatocyte background, in which FXRα1 and α2 were described to differentially affect bile acid and lipid metabolism (Vaquero et al., 2013, Ramos Pittol et al., 2020, Correia et al., 2015, Boesjes et al., 2014). We observed that HCs mostly express FXRα3 and α4, whereas HSCs express mostly FXRα1 and α2. This calls for a careful reexamination of FXR isoforms’ biological properties in each cell (sub)type such as CaHSCs and PaHSCs, which is currently underway in our laboratory.

Taken as a whole, this study provides a compendium of NR expression in parenchymal and non-parenchymal liver cells which calls for an in-depth investigation of NR functions in liver cell populations. In addition to the multiple layers of NR activity regulation, that ranges from ligand availability, dimerization, transcriptional comodulator interaction and post-translational modifications, the expression territory, hence the cellular background is likely to confer specific properties to NR-controlled signaling pathways, and this mapping will provide new guidance for NR-based therapies.

## AUTHOR’s CONTRIBUTIONS

Author’s contributions were: Conceptualization: FPZ, BS, PL; Methodology: FPZ, AB, MV, OMC, LG, CG, DD, MBG, LP; Software: LG, JCD, MJ; Validation: FPZ, OMC, JCD, CG, MJ, LP; Formal analysis: FPZ, AB, MBG, PL; Investigation: FZ, AB, OMC, LP, CG, MBG; Resources: DD, PL, BS; Data Curation: FPZ, LG, JE, PL; Writing: DD, FPZ, PL;

Visualization: FPZ, OMC, PL; Supervision: PL, JE, JCD, DD, BS; Project administration: PL, DD, BS; Funding acquisition: PL, DD, BS

## FIGURE LEGENDS

**Supplemental Figure 1.**
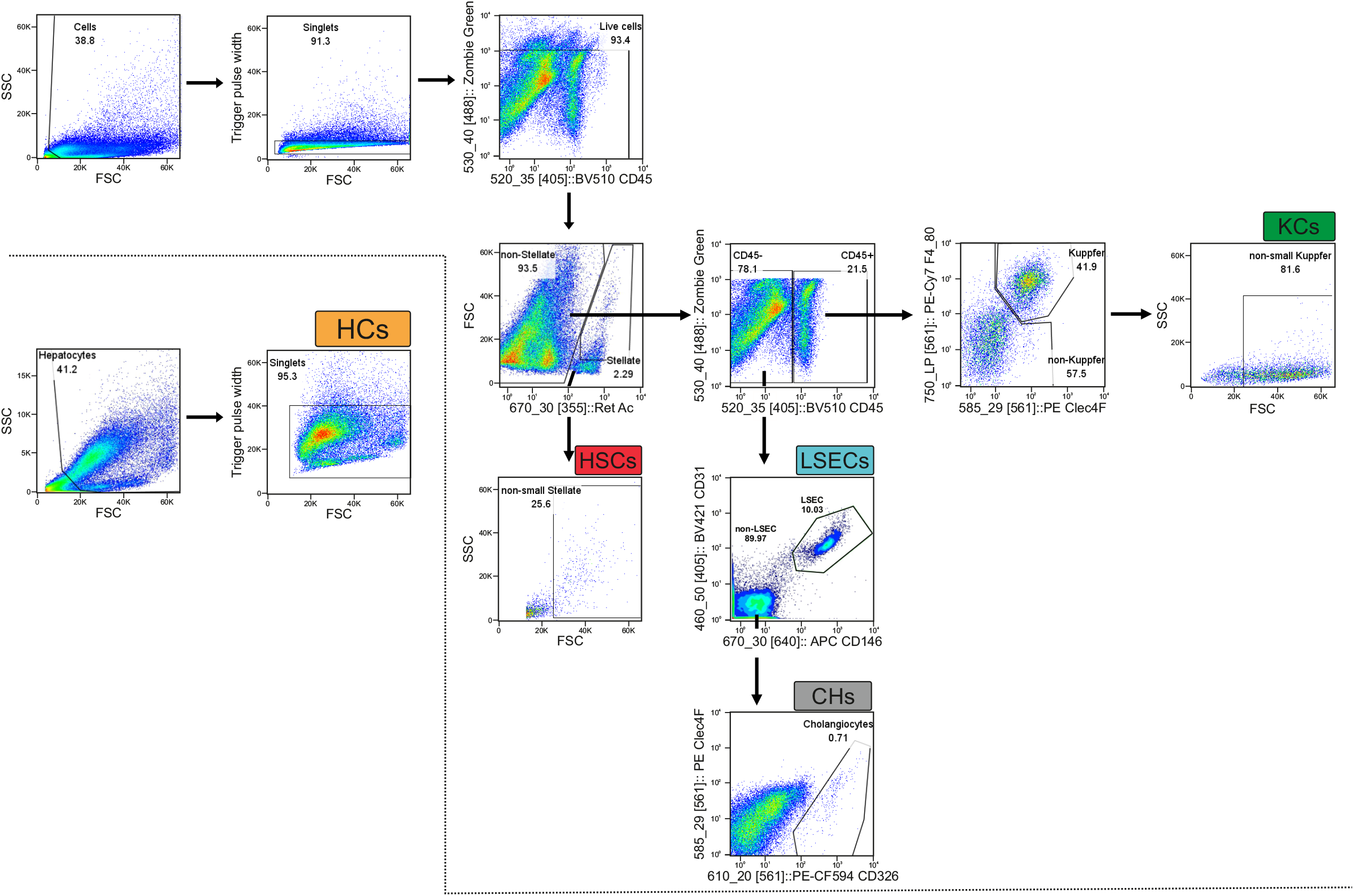
Gating strategy for liver cell type isolation. A representative flow cytometric analysis is shown here. Cells were selected as non-debris on FSC/SSC scatters and singlets were gated on FSC-H/FSC-W. Live cells were selected as “low” for Zombie Green staining. The polychromatic flow cytometry strategy was applied to isolate indicated liver cell types. Further details can be found in the Material & Methods section as well as in Figure 1.

**Supplemental Figure 2.**
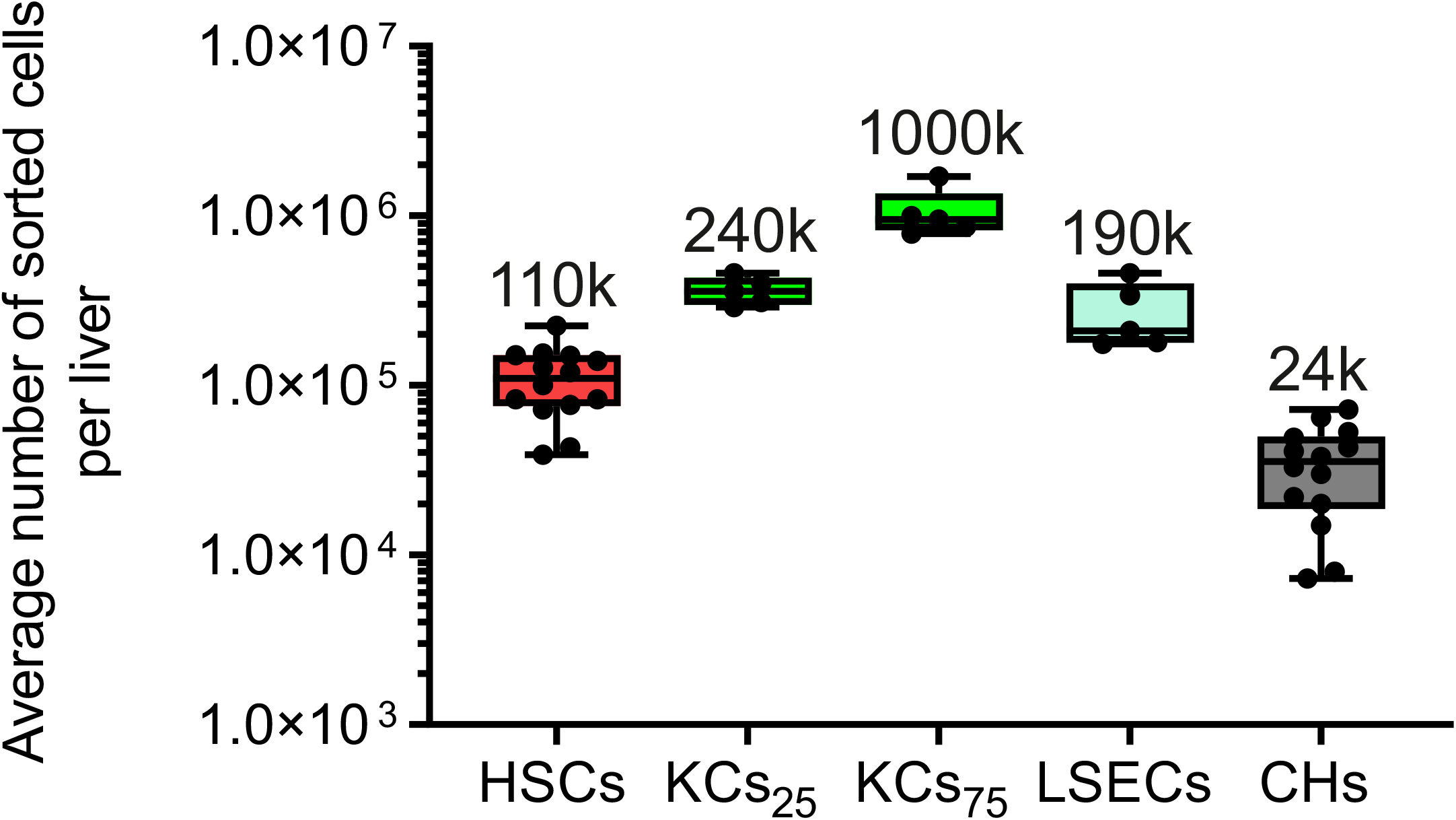
Numbering of purified cells. Cell numbers obtained after the FACS procedure are indicated (n=5-14). KCs Kupffer cells; CHs: cholangiocytes; LSECs: liver sinusoidal endothelial cells; HSCs: hepatic stellate cells.

**Supplemental Figure 3.**
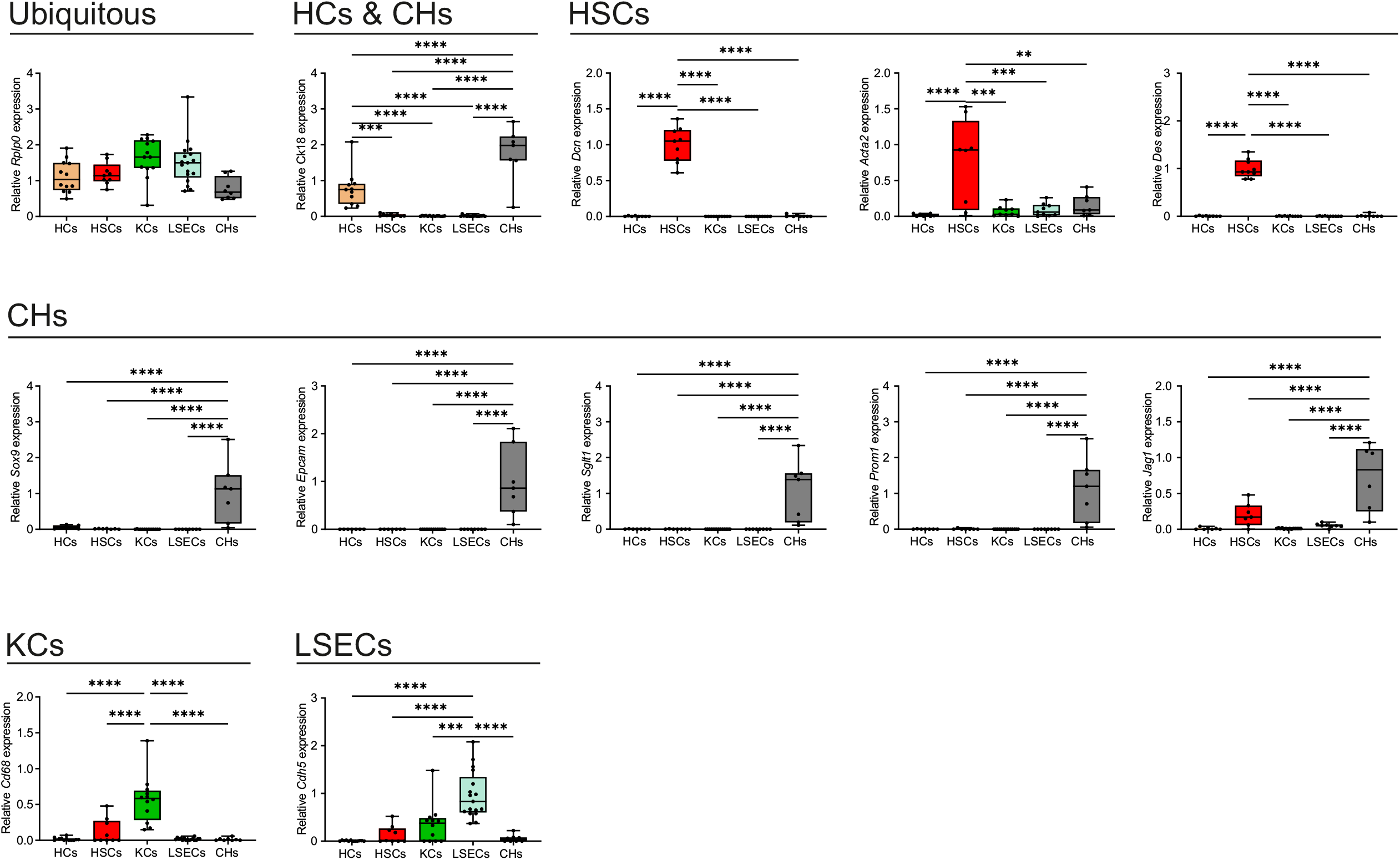
Cell purity assessment by gene expression profiling. RNAs extracted from each purified cell type was analyzed by RT-qPCR (n=9-16) and probed for the expression of ubiquitous (*Rplp0*/*36b4*), common (HCs and CHs, *Ck18*) or specific (HSCs: *Dcn, Acta2, Des*; CHs: *Sox9, Epcam, Sglt1, Prom1, Jag1*; KCs: *Cd68*; LSECs: *Cdh5*) cell markers.

**Supplemental Figure 4.**
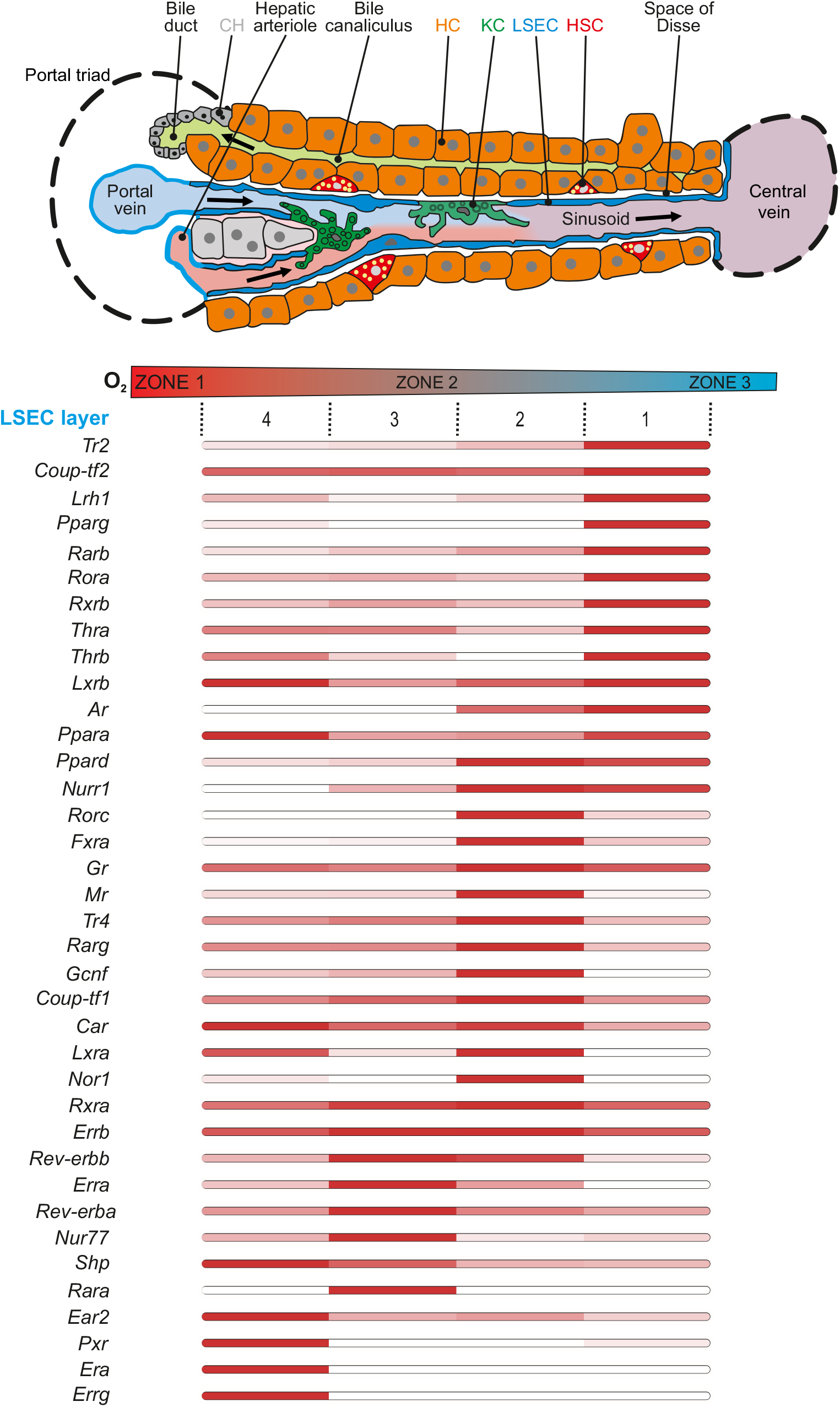
NR zonation in liver sinusoidal endothelial cells. Upper panel: Schematic organization of a liver trabeculae [adapted from Wikimedia Commons and initially published in (Frevert et al., 2005)]. Lower panel: expression values for each NR were extracted from (Halpern et al., 2018) and used to compute an heatmap. Red: High expression, white: low expression. Arrow (right to left) indicates the bile flow, arrows (left to right) indicate the blood flow.

**Supplemental Figure 5.**
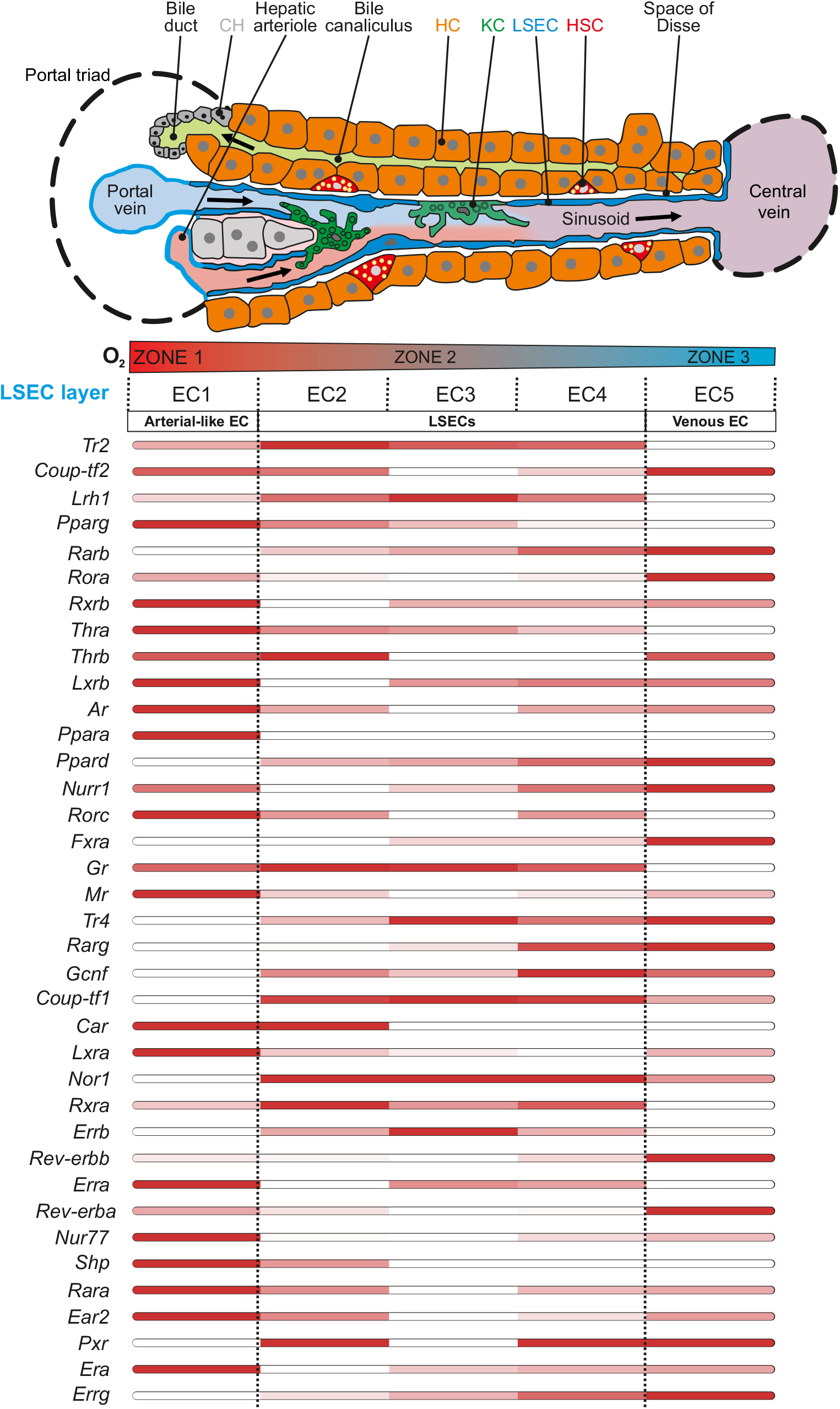
NR zonation in liver sinusoidal endothelial cells. Upper panel: Schematic organization of a liver trabeculae [adapted from Wikimedia Commons and initially published in (Frevert et al., 2005)]. Lower panel: expression values for each NR were extracted from (Su et al., 2021) and used to compute an heatmap. Red: High expression, white: low expression. Arrow (right to left) indicates the bile flow, arrows (left to right) indicate the blood flow.

**Supplemental Figure 6.**
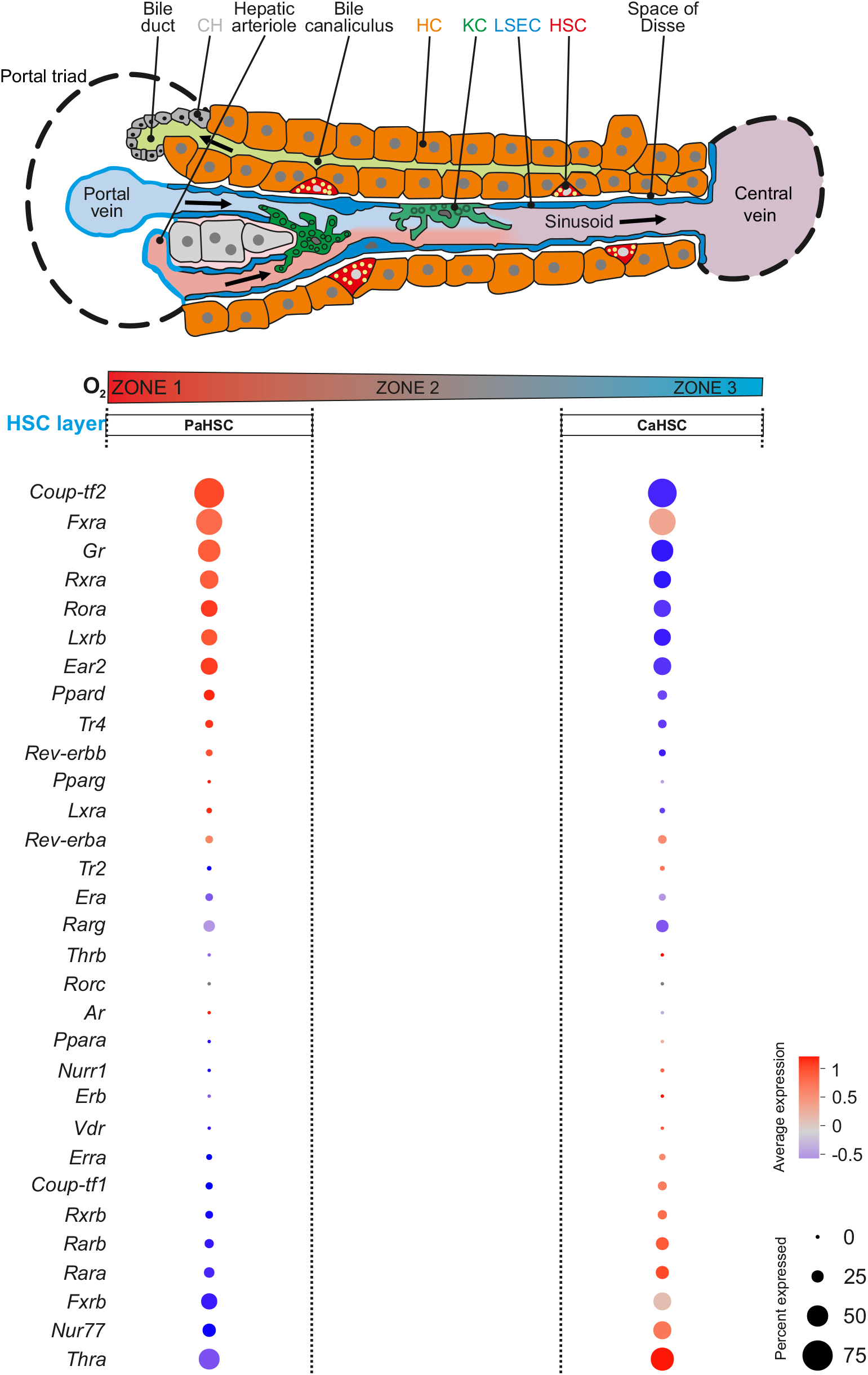
NR zonation in hepatic stellate cells. Upper panel: Schematic organization of a liver trabeculae [adapted from Wikimedia Commons and initially published in (Frevert et al., 2005)]. Lower panel: expression values for each NR were extracted from (Dobie et al., 2019) and used to generate a bubble plot in which the color gradient indicates the expression level (blue: low expression, red: high expression) and the circle diameter indicates the number of cells expressing the transcript. Arrow (right to left) indicates the bile flow, arrows (left to right) indicate the blood flow.

**Table 1. List of primers used in RT-qPCR experiments.**

**Table 2. NR gene counts and NR isoform description.** Sheet 1: Normalized average (n=3) RPKMs are indicated for each mouse NR. The color scale compares NR expression level between cell types (from white, no expression to red, highest expression), numbers are the averaged RPKM.

**Table 3. JTK_cycle output.** Time-dependent transcriptomic data were analyzed using the JTK_cycle R script. Sheet 1: JTK_cycle output for whole liver from ad libitum-fed (AdLib) mice. Sheet 2: JTK_cycle output for whole liver from time-restricted-fed (TRF) mice. Sheet 3: Comparison of the amplitude of 24h-cycling transcripts in “AdLib” and “TRF” conditions.

